# The Relationship Between the Cervical Microbiome, HIV Status, and Pre-Cancerous Lesions

**DOI:** 10.1101/375592

**Authors:** Cameron Klein, Daniela Gonzalez, Crispin Kahesa, Julius Mwaiselage, Nirosh Aluthge, Samodha Fernando, John T. West, Charles Wood, Peter C. Angeletti

## Abstract

Nearly all cervical cancers are causally associated with Human Papillomavirus (HPV). The burden of HPV-associated dysplasias in Sub-Saharan Africa is influenced by HIV. To investigate the role of the bacterial microbiome in cervical dysplasia, cytobrush samples were collected directly from cervical lesions of 144 Tanzanian women. The V4 hypervariable region of the 16S rRNA gene was amplified and deep-sequenced. Alpha diversity metrics; Chao1, PD whole tree, and operational taxonomic Unit (OTU) estimates, displayed significantly higher bacterial richness in HIV positive patients (P = 0.01) than in HIV negative patients. Within HIV positive patients, there was higher bacterial richness in patients with high grade squamous intraepithelial lesions (HSIL; P = 0.13) than those without lesions. The most abundant OTUs associated with high-grade squamous intraepitheilal lesions (HSIL) were *Mycoplasmatales, Pseudomonadales*, and *Staphylococcus*. We suggest that a chronic mycoplasma infection of the cervix can contribute to HPV-dependent dysplasia by sustained inflammatory signals.

## Introduction

Human Papillomavirus (HPV) is the causative agent responsible for 99% of cervical cancers [1]. HPV contributes to about 4.8% of all cancers [1]. The disease burden of HPV is most dramatic in developing regions of the world, with HPV being attributed to 14.2% of cancers in Sub-Saharan Africa [1]. Cervical cancer disproportionately affects Sub-Saharan Africa, where 9% of the world’s female population over 15 years old accounts for 14% of the world’s incident cervical cancer and 18% of cervical cancer-related deaths [2]. The current study uses cervical swab samples obtained from Tanzania, which has among the highest cervical cancer mortality rates by country. Sub-Saharan Affrica also has among the highest HIV rates in the world. The association between HIV and cervical cancer has been better studied than any other factor associated with HPV-related cancers. HIV infection has been strongly linked to increased risk of infection with HPV, and the severity of HPV pathogenesis [3–5]. HR HPV genotypes are more prevalent in HIV+ women, suggesting HIV infection provides an environment where these high-risk HPV’s can better establish infection and replicate [6]. A likely factor in this is a decrease in T-cell surveillance, which results in an increase in HPV replication with decreasing CD4+ cell count, and other changes in the cervical immune microenvironment as HIV infection progresses. Multiple studies have shown an increase in HPV detection, cervical intraepithelial neoplasms in individuals with less than 200 CD4+ cells per μl serum [7–10]. Thus, the cervical immune microenvironment may be a co-factor in suppression of cervical cancer. Changes in the cervicovaginal bacterial microbiome has been suggested to contribute to the development of pre-cancerous cervical lesions [11–17]. Chronic inflammation of the cervix (cervicitis) which is a result of cervicovaginal pathogens, leads to conditions like Pelvic Inflammatory Disease (PID) and Bacterial Vaginosis (BV), both of which are associated with persistent HPV infection and cervical cancer [18, 19]. Both PID and BV are more prevalent in Sub-Saharan Africa and in HIV positive populations [20–22]. Comparative genomic analyses in women infected with HIV have shown that a shift in microbial diversity as a result of BV is detectable; if this shift directly effects formation of pre-cancerous cervical lesions is not clear [23]. Given that cervical cancer rates which are expected to rise in Sub-Saharan Africa as the HIV positive population receives life-extending ART, it is even more important to understand the risk factors associated with the cervical microbiome. There are currently no studies that define how cervical microbiota differ at different stages of cervical cytology or as a function of HIV status. The current study defines bacterial communities associated with cervical lesions and with HIV, which represents a significant advance. Cervical cytology is graded by pap smear screening for nuclear abnormalities. There are five stages: negative for intraepithelial lesion or malignancy (NILM), atypical squamous cells of undetermined significance (ASC-US), low-grade squamous intraepithelial lesions (LSIL), atypical squamous cells: cannot exclude high-grade lesions (ASC-H), and high-grade squamous intraepithelial lesions (HSIL). In this study, we utilized 16s rRNA gene deep-sequencing on a set of 144 cervical swab samples from a cohort of Tanzanian women to gain an understanding of the differences in the cervical bacterial community composition as a function of cervical cytology grade and HIV status. The data presented here identifies bacterial taxonomies associated with high-grade cervical lesions with a new level of precision since swab samples samples were take directly from cervical lesions. The rationale behind this approach was that the sight of the lesion is where tumors form, thus it bacteria associated at this sight were more likely to be relevant to disease status.

## Methods

### Participants and Ethical Precautions

This study reports findings derived from a larger cross-sectional cohort study analyzing demographics of HPV and cervical cancer in HIV positive and negative women from rural and urban Tanzania. All human subjects protocols were approved by safety committees at Ocean Road Cancer Institute (ORCI) and the University of Nebraska-Lincoln in accordance with the Helsinki Declaration. Participation by patients was entirely voluntary and written patient consent was required for inclusion in the study.

Tanzania is part of an ongoing study site to follow HIV and associated secondary viral infections in women at ORCI, the only cancer treatment hospital in Tanzania. Between March 2015 and February 2016, female patients undergoing cervical cancer screening were approached for enrollment in the study. Those who were pregnant, menstruating, under 18, reported being sick in the past 30 days, or had a preexisting, non-HIV, immunologic defect were excluded from the study. Disease histories as well as physical examinations were carried out to rule out any clinical symptoms or visible signs for these conditions. Samples were collected at three sites: ORCI in Dar es Salaam, and rural clinics in Chalinze and Bagamoyo. A total of 144 cervical cytobrush samples obtained from these women were sequenced, of which 138 produced at least 1000 reads, and 132 included successful measurements of both HIV status and cervical cytology.

### Demographic Data Collection

All study participants who gave informed consent and were evaluated by study clinicians. A set of pre-tested, standardized questionnaires were used to gather demographic data. All personal identifiers were removed from samples to ensure patient confidentiality. With patients’ permission, medical history was retrospectively retrieved from hospital medical records. Over 30 variables were identified and assessed in the questionnaire. The current study only uses data collected regarding age and laboratory test results (pap smears, VIA, CD4 count, genotyping of HPV, results of serological testing for HIV-1).

### Sample Collection

Blood specimens were collected via venipuncture into acid-citrate-dextrose tubes and processed using centrifugation at the on-site study laboratory within 6 hours of being drawn. The separated plasma was tested at ORCI using Standard Diagnostics HIV-1/2 3.0 detection kit and BD products CD4 FITC, CD8 PE, and CD3 PerCP antibodies to test CD4 count on a BD Accuri C6 Plus. Cervical cytobrush samples and pap smears were collected from all patients. Pap smears were examined by at least three trained cytologists, and classified according to the pap classification protocol: negative for intraepithelial lesion or malignancy (NILM), atypical squamous cells of undetermined significance (ASC-US), low-grade squamous intraepithelial lesions (LSIL), atypical squamous cells: cannot exclude high-grade lesions (ASC-H), and high-grade squamous intraepithelial lesions (HSIL). Cervical cytobrush specimens were placed in lysis buffer, then shipped to the Nebraska Center for Virology at the University of Nebraska-Lincoln (UNL) for processing.

### DNA isolation, 16S rRNA Library Preparation, and Sequencing of the V4 Region

Cervical cytobrush samples were vortexed and separated from the brush with lysis buffer. DNA was extracted from the lysis buffer using the Qiagen Tissue extraction kit (Dneasy) according to the manufacturer’s protocol. The DNA concentration was determined by UV spectrophotometer at 260/280 nm.

DNA was then used for tag sequencing of the V4 hypervariable region of the 16S rRNA gene. A 250 base pair section of the V4 region was amplified using universal primers described in [24]. The PCR reactions were performed in 25μL. The cycling conditions were an initial denaturation of 98°C for 3 minutes, followed by 25 cycles of denaturation at 98°C for 30 seconds, annealing at 55°C for 30 seconds, and extension at 68°C for 45 seconds, then a final elongation of 68°C for 4 minutes. Following amplification, PCR products were analyzed on a 2% agarose gel to confirm correct product size. Normalized amplicons (1-2 ng/μL) from 144 samples were pooled together using an epMotion M5073 liquid handler (Eppendorf AG, Hamburg, Germany). Pooled libraries were sequenced using the Illumina MiSeq platform using the dual-index sequencing strategy outlined by Kozich et al. [25].

### Data Processing and Bacterial Community Analysis

The sequencing data obtained from the sequencer was subsequently analyzed using the Illumina MiSeq data analysis pipeline used by the Fernando lab (described in detail in: https://github.com/FernandoLab). Briefly, initial quality filtering was carried out to remove sequences which had ambiguous bases, incorrect lengths, and inaccurate assemblies. Subsequently, the quality filtered reads were run through the UPARSE pipeline (http://www.drive5.com/uparse/) and subjected to chimera filtering and OTU clustering (at a similarity threshold of 97%), followed by the generation of an OTU table. Taxonomy was assigned to the OTUs using the assign_taxonomy.py command available in QIIME using the latest version of the Greengenes database (May 2013).

### Statistical Analyses

The OTU table was rarefied across samples to the lowest sample depth (1000 reads) using QIIME based on the Mersenne Twister pseudorandom number generator. All statistical analyses were performed with samples at an even depth. Bar charts summarizing average taxonomic makeup of samples by HIV status and cervical cytology were constructed from the rarefied OTU table in QIIME. Heatmaps showing the relative abundance of bacterial taxonomic families were constructed using the ‘plot_ts_heatmap’ command using the mctoolsR package for R. Differences in bacterial families by HIV status or cervical cytology were evaluated using the ‘taxa_summary_by_sample_type’ command in mctoolsR using Kruskal Wallace. Families with less than 1% abundance were excluded in this analysis. Alpha diversity estimators Chao1, observed OTUs, and PD whole tree and rarefaction curves were calculated for the overall bacterial community using QIIME. Good’s coverage test was performed to evaluate if adequate sampling depth was achieved. Mean alpha diversity estimates for HIV positive, HIV negative, NILM, LSIL, and HSIL groups were compared using nonparametric two-sample t-tests using Monte Carlo permutations in QIIME. The weighted and unweighted UniFrac distance matrix for bacterial communities were calculated using QIIME. Even depth across samples avoided biases that could be encountered when using the Unifrac metric [26]. Bacterial community composition differences were evaluated using the unweighted UniFrac distance matrix as an input for a distance-based redundancy analysis (db-RDA) in Qiime, where HIV status, cervical cytology, or HPV status were used as main effects. A heatmap was generated using the heatmap.2 command in the “ggplots” package for “R” using the Bray-Curtis distance matrix to visualize relationships between samples. Significance was declared at P ≤ 0.1 throughout this study. The linear discriminant analysis effect size (LEfSe) was used to identify specific OTUs that differed HIV status and cervical cytology [27]. LEfSe uses a non-parametric factorial Kruskal-Wallis sum-rank test followed by a linear discriminate analysis to identify both statistically significant and biological relevant features. The OTU relative abundances were used as an input for LEfSe. Demographic data was examined using odds ratio and an associated p value to test for factors associated with HIV status and/or a positive VIA status. All p values are reported as FDR corrected p values.

### Ethics Statement

All human subjects protocols were approved by safety committees at Ocean Road Cancer Institute (ORCI) and UNL in accordance with the Helsinki Declaration. Participation by patients was entirely voluntary and written patient consent was required for inclusion in the study.

## Results

### Demographics

Of the 144 patient samples sequenced, 41 were HIV+ and 103 HIV−, with an average age of 37 years old. Cervical cytology was measured to be NILM in 23 samples, LSIL in 72 samples, and HSIL in 50 samples. Visual inspection with acetic acid (VIA), the standard for cervical lesion detection in Tanzania, was carried out immediately following sample collection. Twenty-six patients were found to be VIA positive for cervical lesions and 115 VIA negative. All VIA positive samples were identified as LSIL or HSIL, while several VIA negative samples were found to be NILM, LSIL or HSIL by pap smear. Odds ratios were used to identify risk factors for testing VIA positive. Testing HIV+, HSIL, having >5 sexual partners, and having been infected with a sexually transmitted infection (STI), were identified as risk factors for positive VIA status (p=0.0001, p=0.038, p=0.006, p=0.0008 respectively).

### Cervical Bacteria Composition and Richness

Samples rarefied to an even depth (1000 reads) were used to generate 813 OTUs. To assess if the sampling depth was adequate, rarefaction curves were generated using observed OTUs for HIV status and cervical cytology (Figure S1). Rarefaction curves for both did not converge, but showed a diminishing rate of new OTU identification as the number of reads per sample increased, implying that sampling depth was adequate for evaluating dominant members of the cervical bacterial community. The Good’s coverage test showed the sequencing depth was able to characterize 99.4% of the bacterial community on average.

The taxonomic analysis of the reads revealed the presence of 6 main phyla (relative abundance >1%) in the cervical epithelium, regardless of HIV or cervical cytology status (Figure 1). Firmicutes were the predominant phylum across all sampling groups, accounting for 41.3% of total reads. The average relative abundance of Firmicutes decreased slightly in HIV+ samples compared to HIV− samples (44.4% to 40.2%), and varied by cervical cytology, though no obvious trend was apparent. When considering only the HIV+ samples, Firmicute relative abundance appeared to decrease in patients with cervical lesions. Firmicute reads were primarily from the genus Lactobacillus, which accounted for 21.9% of total reads. Tenericutes accounted for 1.5% of total reads, and showed a clear increase in relative abundance with increasing severity of cervical lesions. In HIV− patients, tenericutes increased from 0.3% of reads in NILM patients, to 1.3% in HSIL. In HIV+ patients the shift is larger; Tenericute relative abundance increased from 0.2% in NILM patients to 5.0% in HSIL patients. Tenericute reads were primarily assigned to the genus Mycoplasma and Ureaplasma, which account for 1.1% and 0.2% of total reads respectively. Proteobacteria, Fusobacteria, Bacteroidetes, and Actinobacteria had smaller, or less consistent shifts in relative abundance between HIV and cervical cytology categories. The relative abundance of Tenericues and Bacteroidetes were significantly different between HIV+ and HIV− groups (p=0.020, p=0.017 respectively). No other phyla reached significance based on HIV status or cervical cytology. Comparison of relative abundance of bacterial families (Figure 2) found that Mycoplasmataceae and Prevotellaceae were significantly more abundant in HIV+ patients (p=0.03, p=0.07 respectively). No families were found to be significantly different in abundance based on cervical cytology. When separated by HIV status, the differences between bacterial families remained insignificant in HIV− patients, however in HIV+ patients, Prevotellaceae was found to be significantly more abundant in cervical lesions (p=0.068).

**Figure 1:**
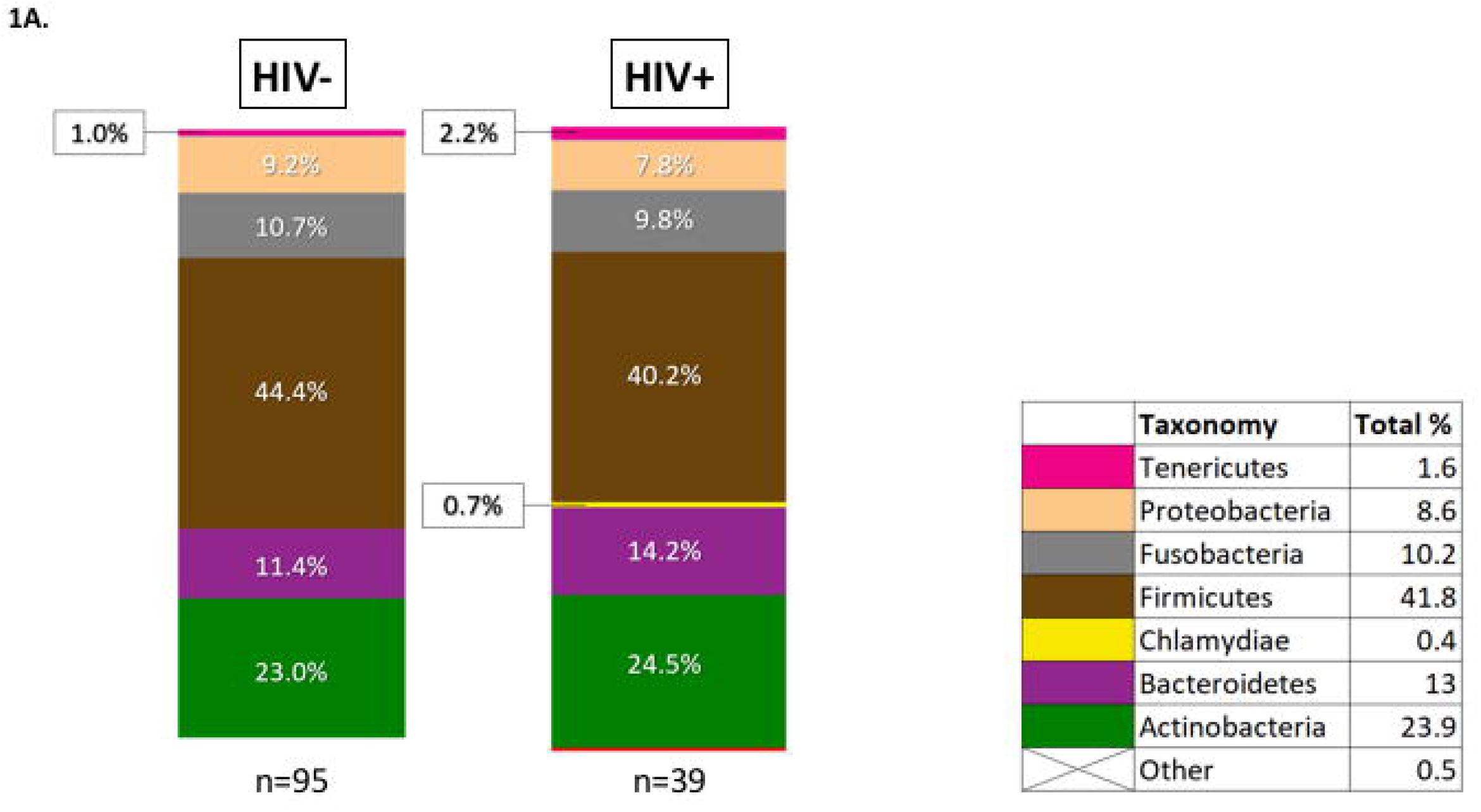
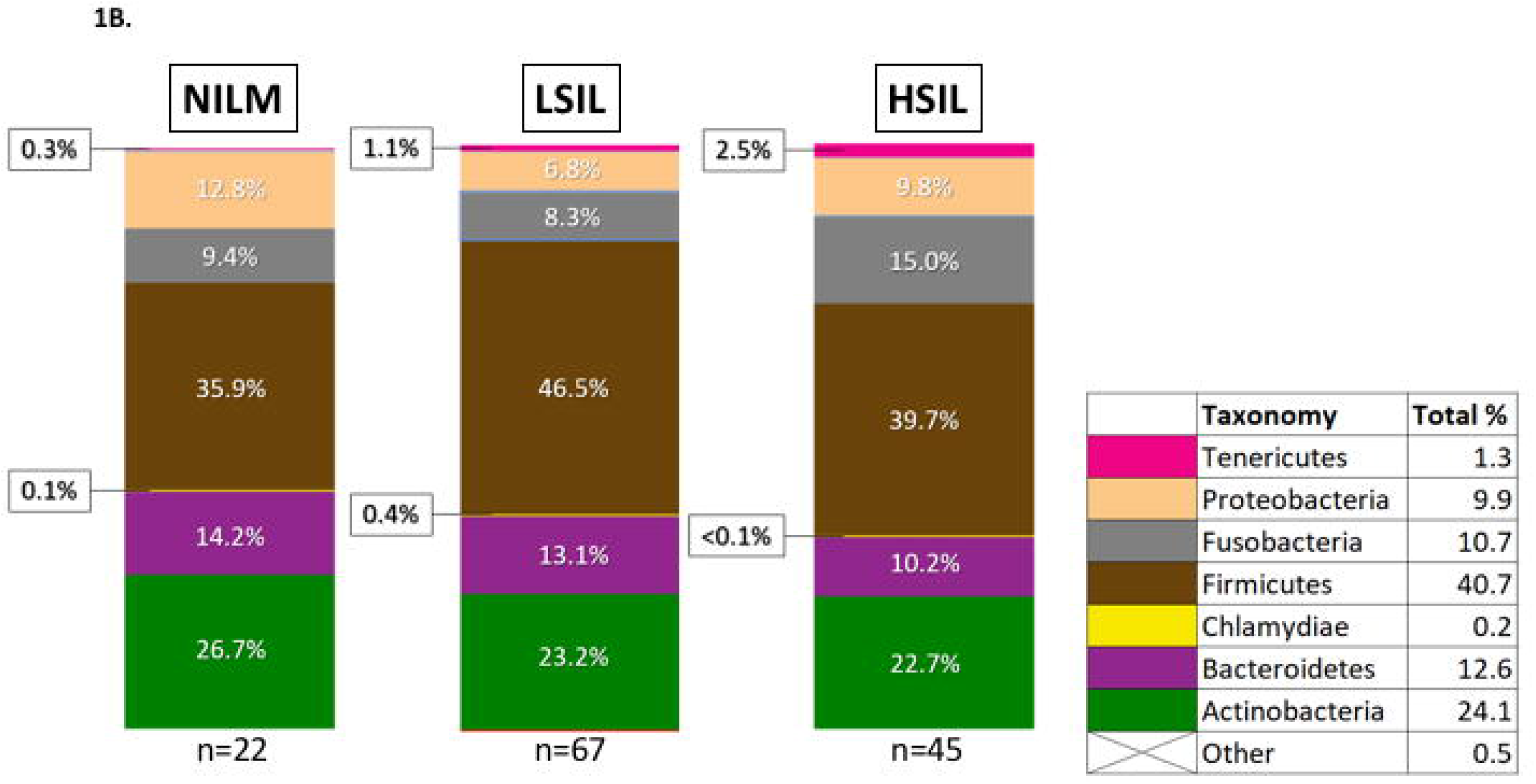
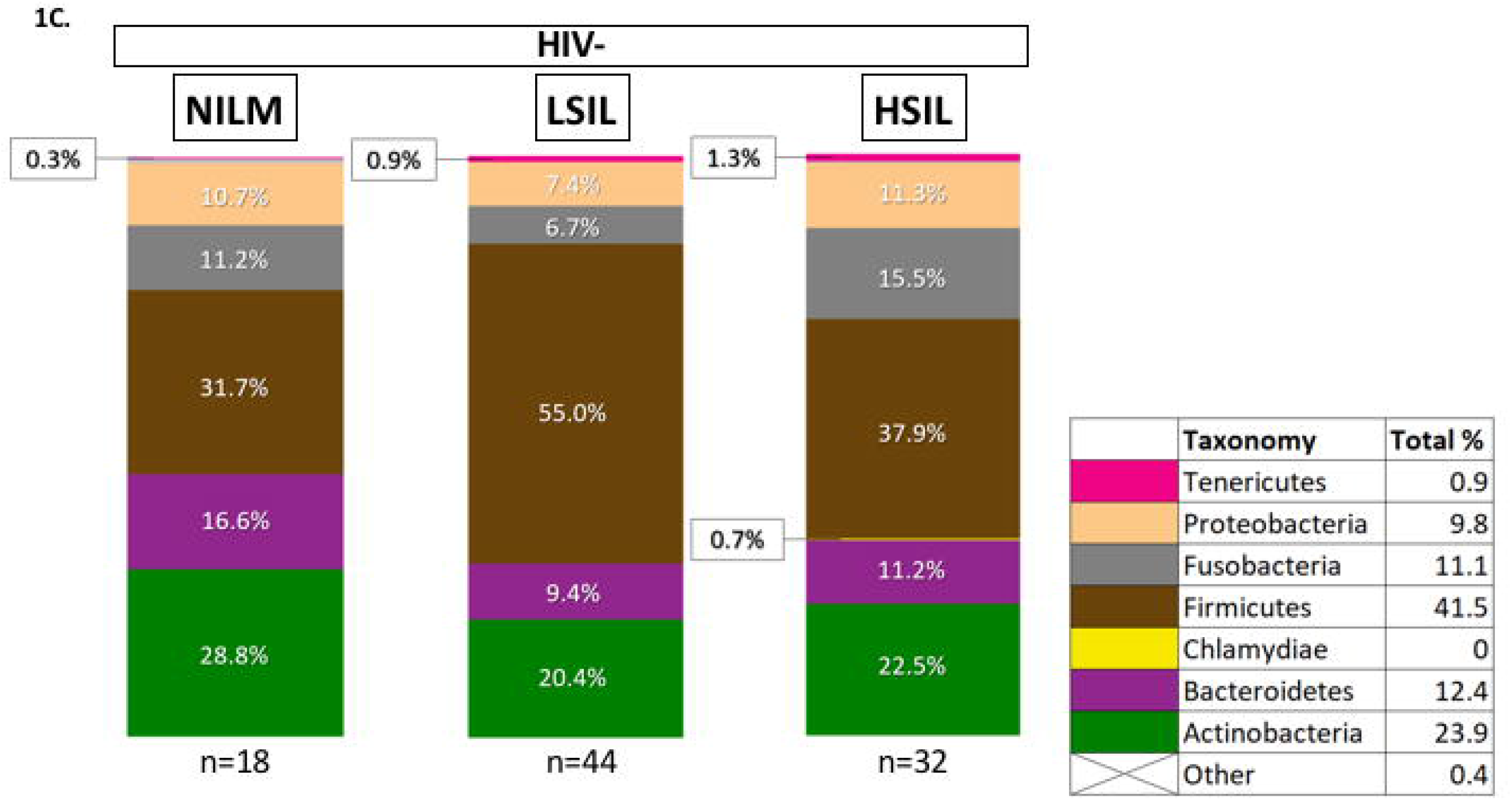
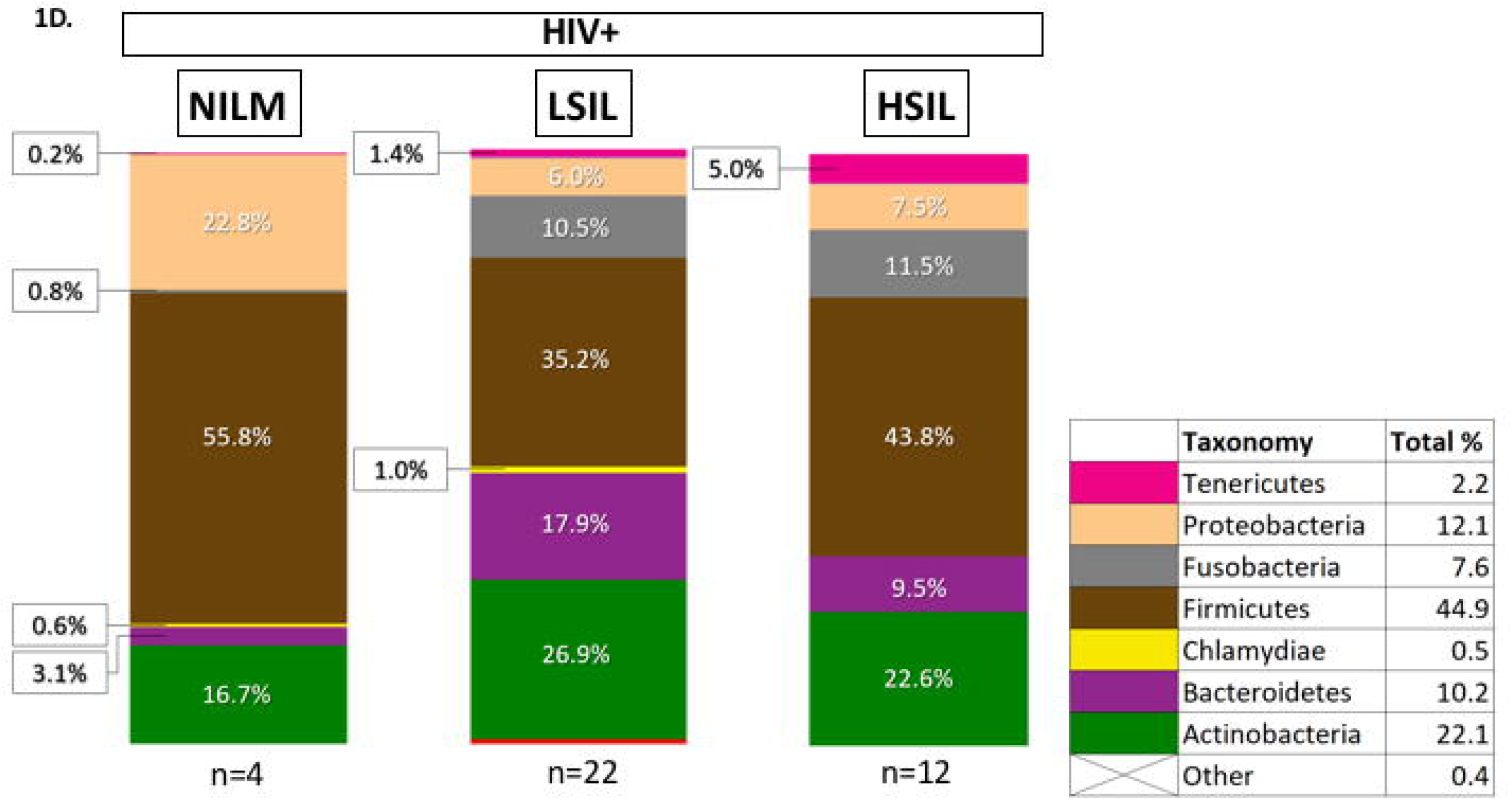
Phylum-level taxonomy of the cervical bacterial community composition as a function of HIV status and cervical cytology. **A**. Phylum-level bacterial taxonomy of the cohort is displayed by HIV status. **B**. Phylum-level bacterial taxonomy of the cohort is displayed as a function of cervical cytology. **C**. Phylum-level taxonomy of HIV negative patients as a function of cervical cytology grade. **D**. Phylum-level taxonomy of HIV posative patients as a function of cervical cytology grade. Phyla are represented as a percent total.

**Figure 2:**
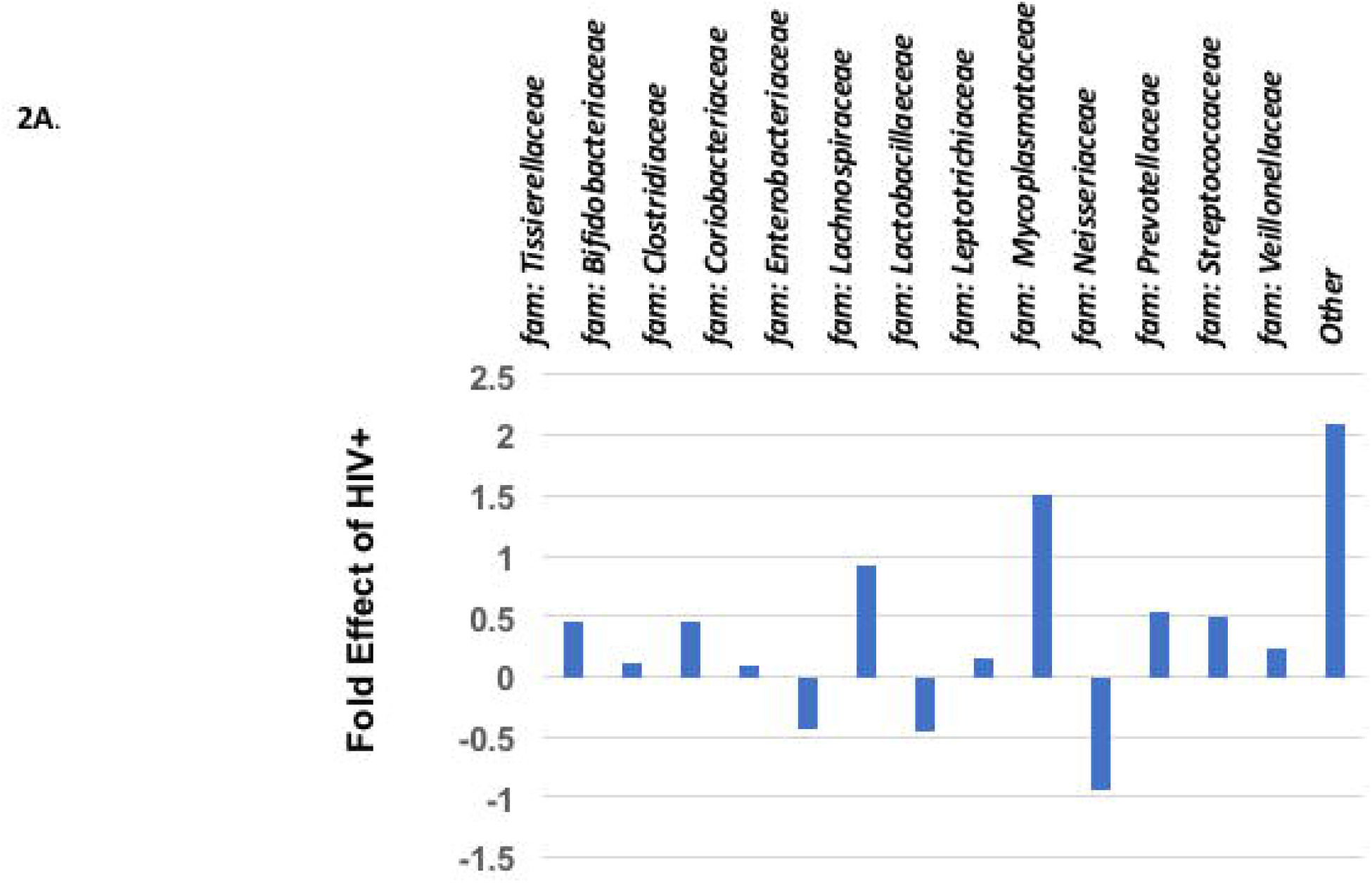
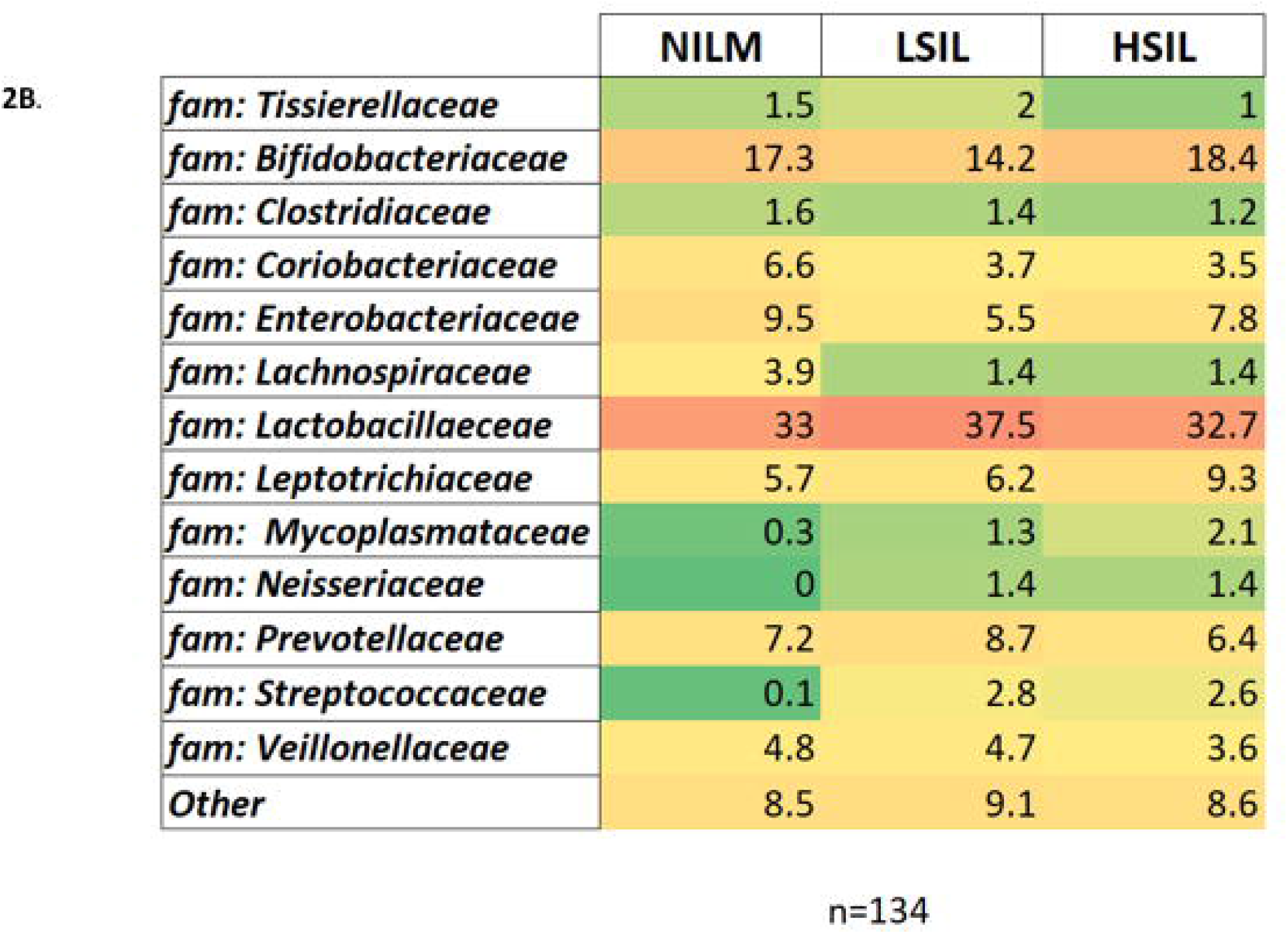
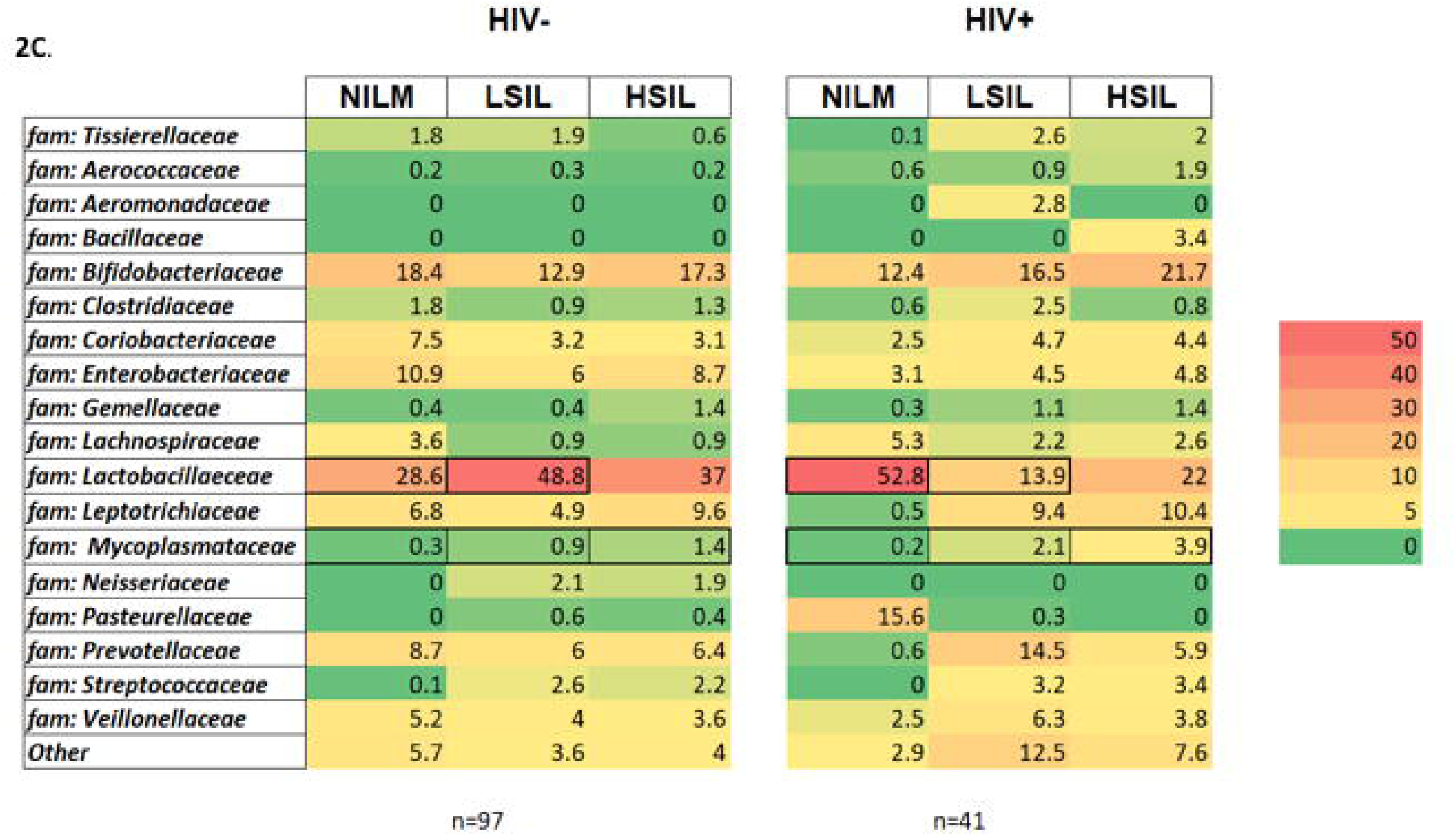
Relative abundance heatmap of family-level taxonomy of the cervical bacterial community composition as a function of HIV status and Cervical Cytology. **A**. Fold effect of HIV+ on the family-level bacterial taxonomy within the cohort. **B**. Relative abundance heatmap of the family-level taxonomy of cohort versus cervical cytology **C**. Relative abundance heatmap of the family-level taxonomy of cohort by cervical cytology, separated by HIV status. The data are presented as percent total. The scale is shown on the right.

### Cervical Bacterial Diversity Estimates

Alpha diversity metrics, Chao1, observed OTUs, and PD Whole Tree, displayed higher (p = 0.009) bacterial richness in HIV+ compared to HIV− patients (Figure 3). A subset of these samples were matched such that the HIV− and HIV+ groups consisted of the same number of samples, with the same average age, and the same contribution of each cervical cytology to help to control for effects of these confounding variables, and ensure differences in diversity estimates are not due to differences in sample size. In this matched subset, estimates also displayed higher (p = 0.003) bacterial richness in HIV+ patients.

**Figure 3:**
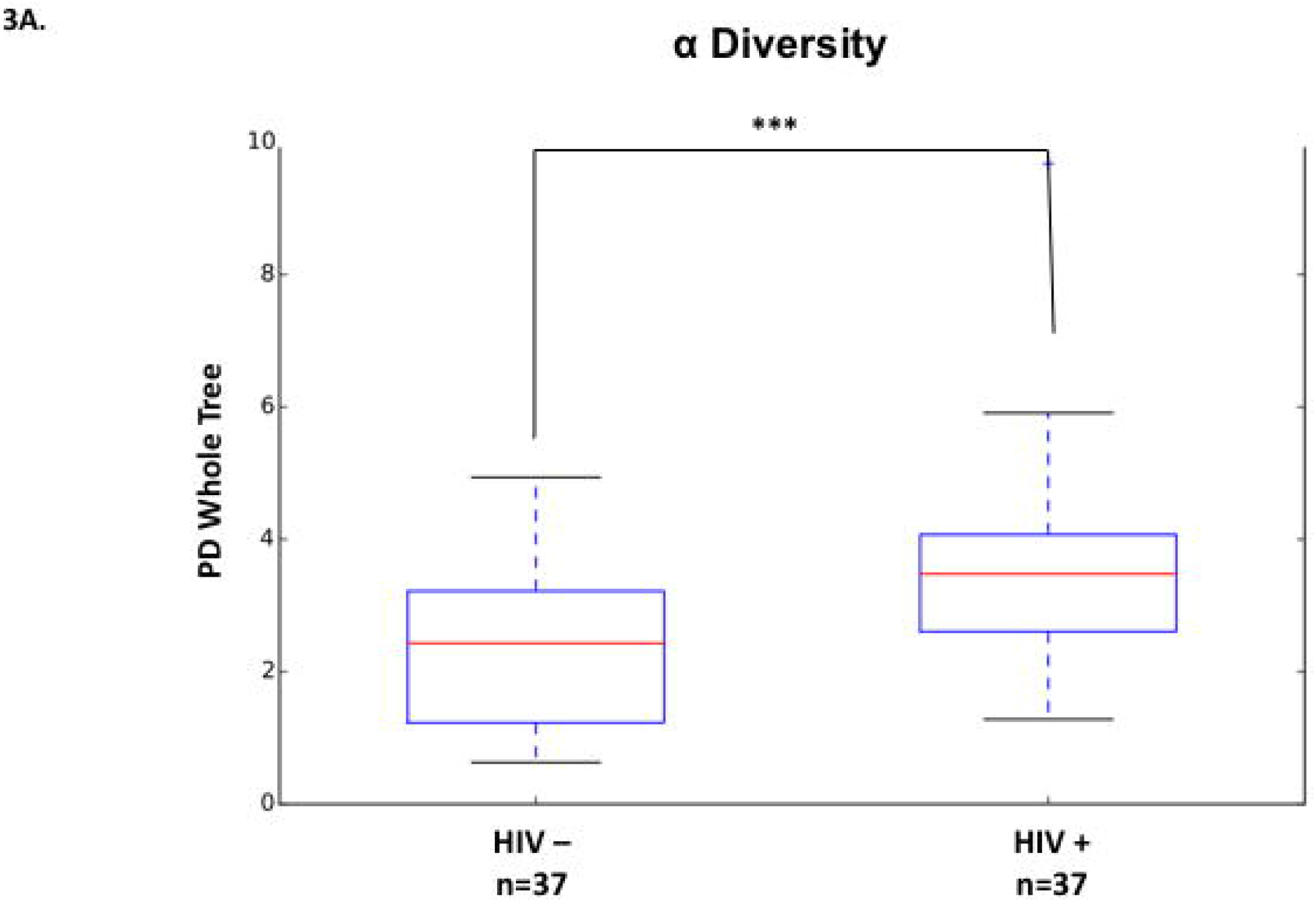
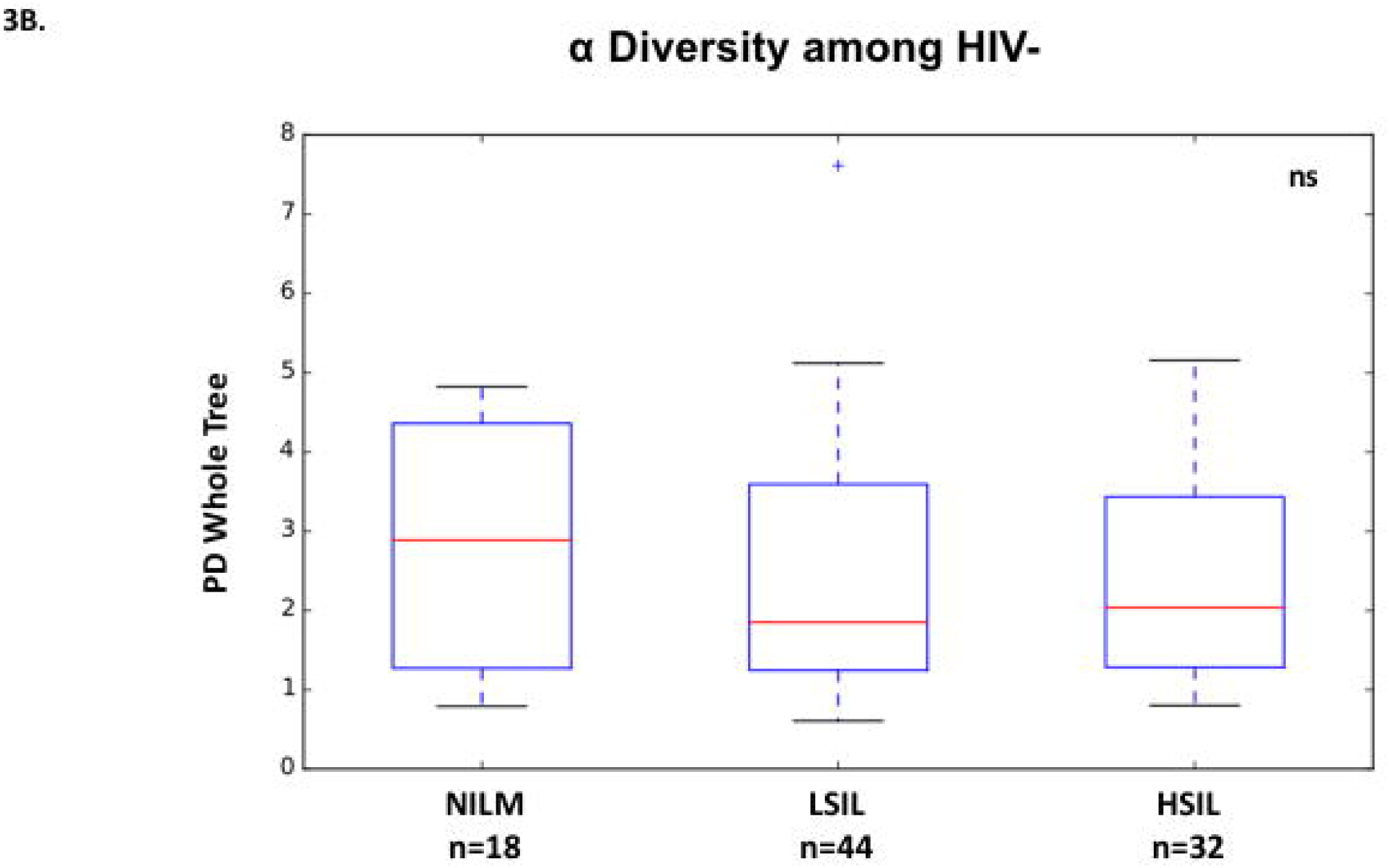
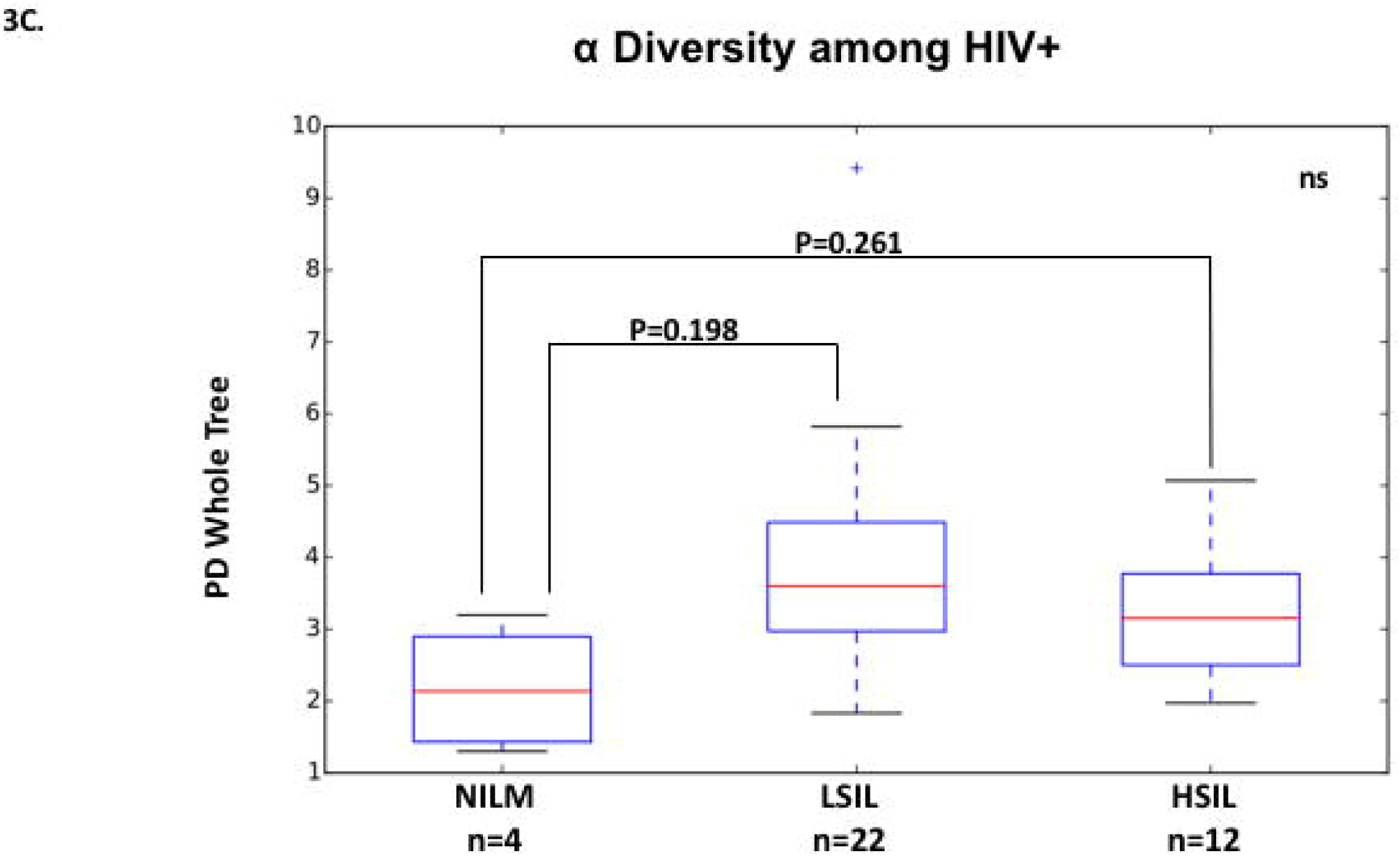
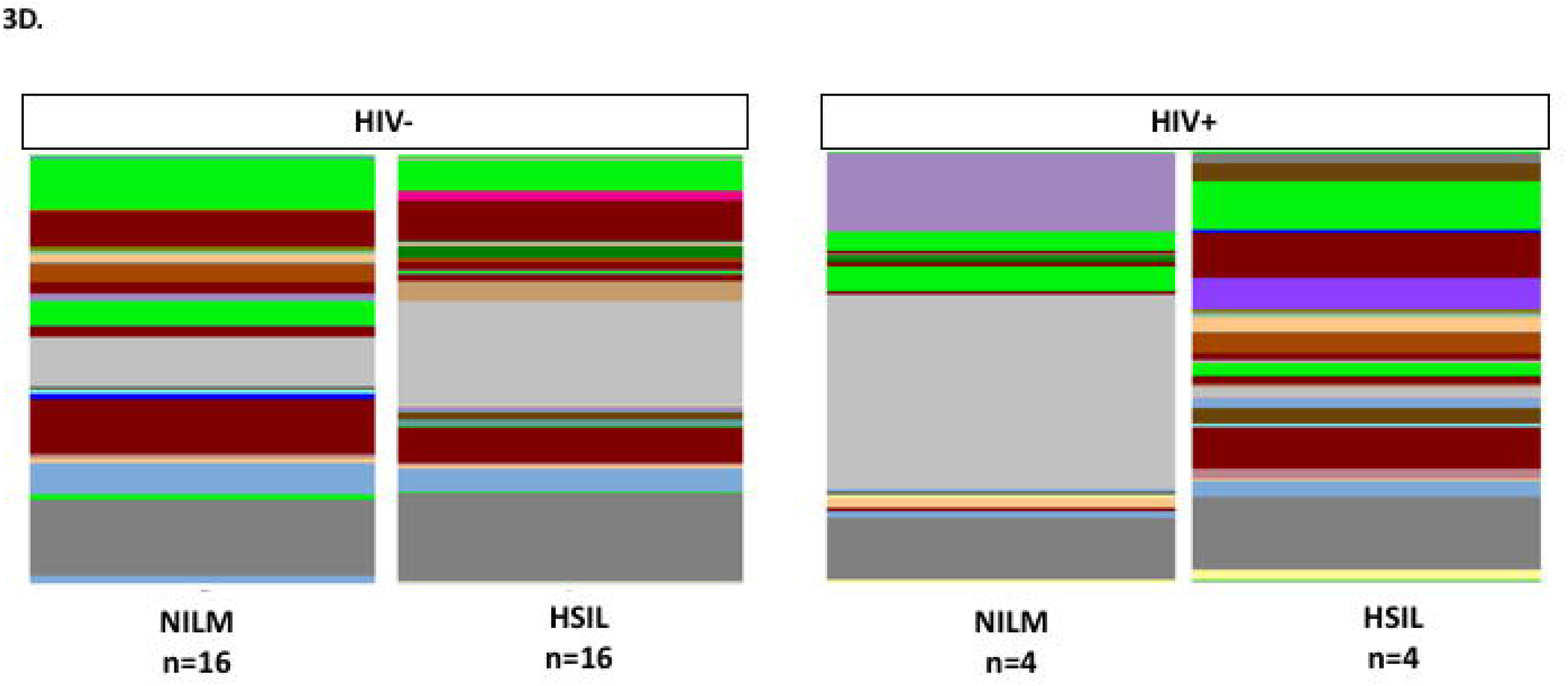
Alpha diversity measurements of cohort sub-groups. **A**. Relationship between HIV status and alpha diversity of cervical bacteria. **B**. Relationship between cervical cytology and bacteria alpha diversity, separated among HIV− individuals. **C**. Relative abundance of genus-level reads differentiated by cervical cytology in HIV+ and HIV− (ns = non-significant, * p < 0.1, ** p < 0.05, *** p < 0.01).

Alpha diversity metrics were similar (P > 0.50) for the samples from patients at different cervical cytology grades (NILM, LSIL, or HSIL) in both matched and unmatched sets. When alpha diversity metrics were compared between cervical cytology groups separately for HIV+ samples, LSIL and HSIL trended towards a higher diversity compared to NILM (p=0.198, p=0.261 respectively). Analysis of age matched, HIV+ NILM/HSIL pairs maintained this trend (p=0.264, Chao1 p=0.13). Comparison of the relative abundance of Genus-level reads between these groups showed a noticeably more diverse profile for HSIL samples, which lack the dominance of Lactobacillus and Haemophilus seen in NILM samples.

Beta diversity analysis showed that bacterial communities were quite varied between samples (Figure 4), no discreet communities characterized a large number of samples. On average, HIV positive patients were shown to have significantly different cervical bacterial communities from HIV negative patients (p=0.001). Similarly, patients who tested positive for HPV tended to have different bacterial communities from those who tested negative (p=0.008). Bacterial communities were also shown to differ significantly depending on cervical cytology among HIV positive patients (p=0.05).

**Figure 4:**
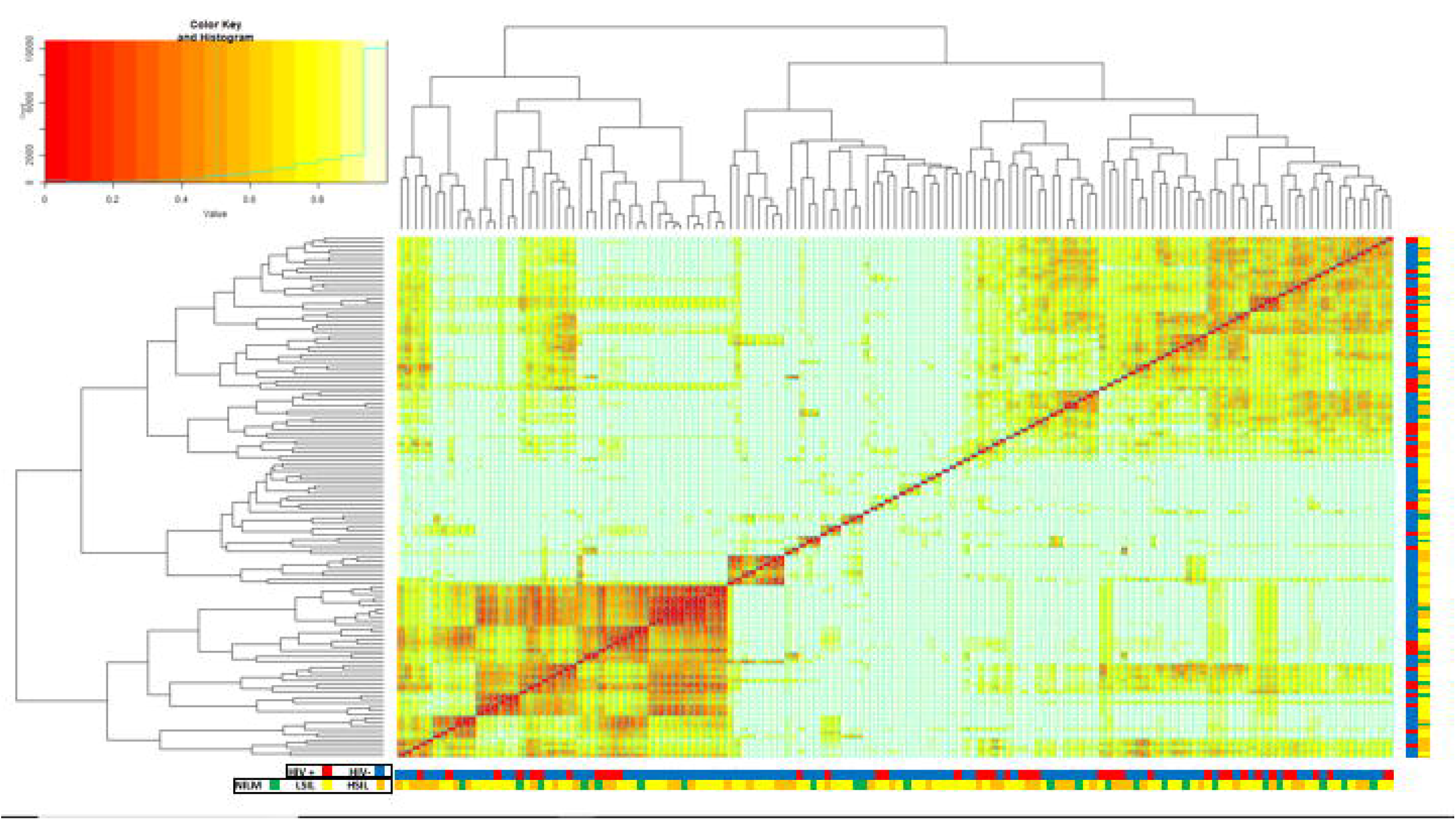
Heatmap of the Bray-Curtis distances between each sample (Beta-Diversity). Samples are grouped into a similarity tree based on abundance of each OTU. Lower values (red) indicate more similarity. HIV status and cervical cytology of each sample is labeled by color beneath each column and beside each row (NILM= Green, LSIL= Yellow, HSIL= Orange).

### Bacteria Associated with Cervical Cytology States and/or HIV Status

Linear discriminant analysis Effect Size (LEfSe) was used to identify bacterial taxonomies which differentiate cervical microbiota in normal individuals (NILM) from microbiota in patients with pre-cancerous lesions (HSIL). The sum of reads at each taxonomic rank was considered. Gammaproteobacteria, s24_7, Paraprevotellaceae (non-verified taxonomy), and Finegoldia associated with NILM cervices, while Pseudomoriadaceae, Staphylococcus, and Mycoplasmatales associated with pre-cancerous lesions. Mcoplasmatales were dominant among Tenericutes, resulting in the significant association seen between the phylum and cervical lesions. LEfSe was then used to compare HIV+, age-matched pairs of NILM and HSIL patients to determine which bacteria may influence the development of lesion in high-risk, HIV+ populations. Mycoplasmatales were most strongly associated with cervical lesions in HIV+ paitents, followed by Parvimonas and Streptococcus. In NILM patients, an abundance of Lactobacillus, especially *L. iners* was found, and somewhat less significantly Finegoldia. LEfSe analysis of samples by HIV found several bacteria to be associated with being HIV+ (Figure 5c). An abundance of non-Lactobacillus Bacilli was the most significant differentiating taxonomy between HIV positive and negative samples. Mycoplasma was also associated with HIV+ individuals, supporting the significant difference in relative abundance between HIV positive and negative groups shown previously using a direct Kruskal Wallace comparison. Interestingly, Ureaplasma (a member of Mycoplasmatales) and *Lactobacillus reuteri* were associated with HIV− patients, while other members of their respective families were associated with HIV+ patients. This suggests the existence of metabolic niches in the cervical microbiome which may be populated by pathogenic or non-pathogenic associating bacteria.

**Figure 5:**
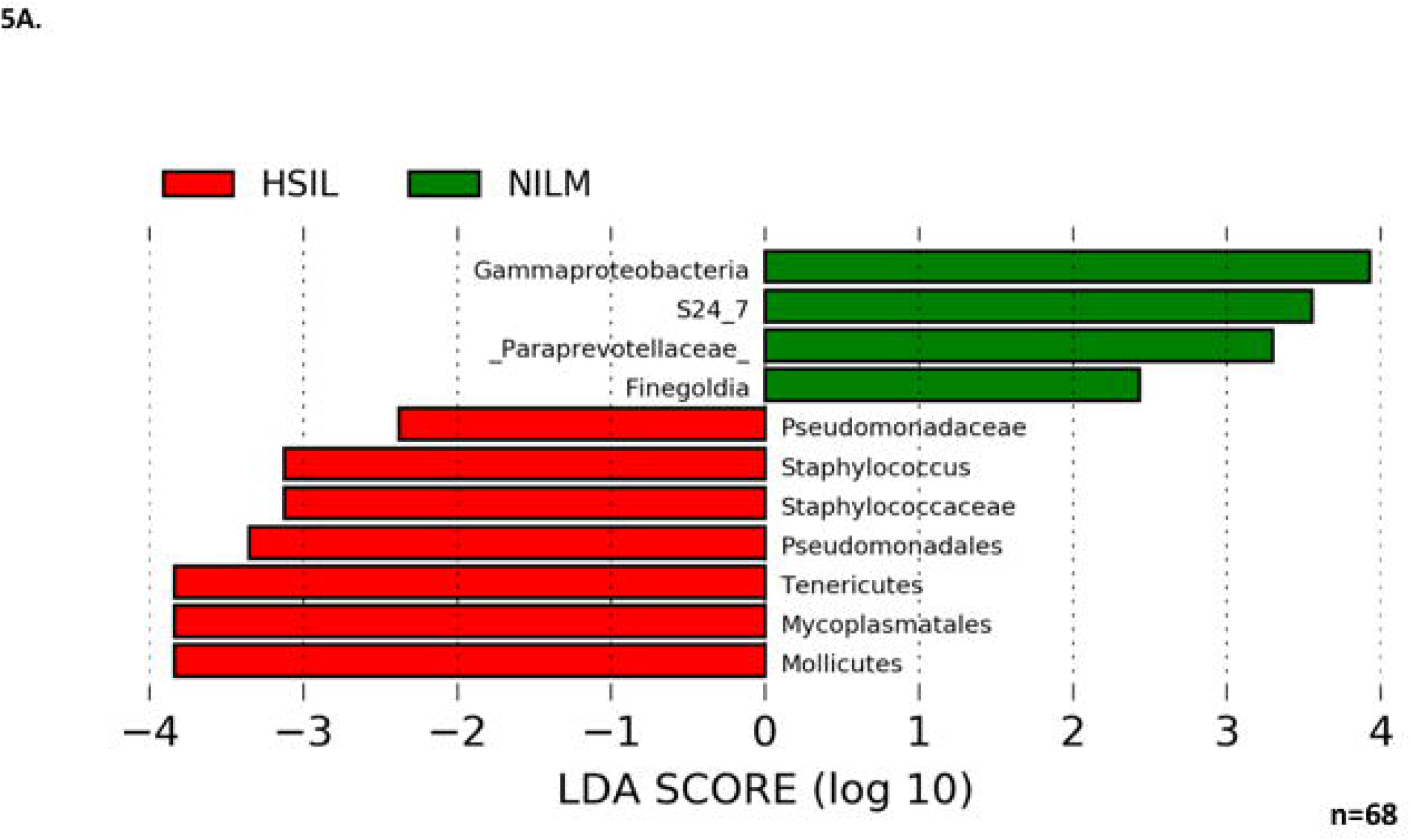
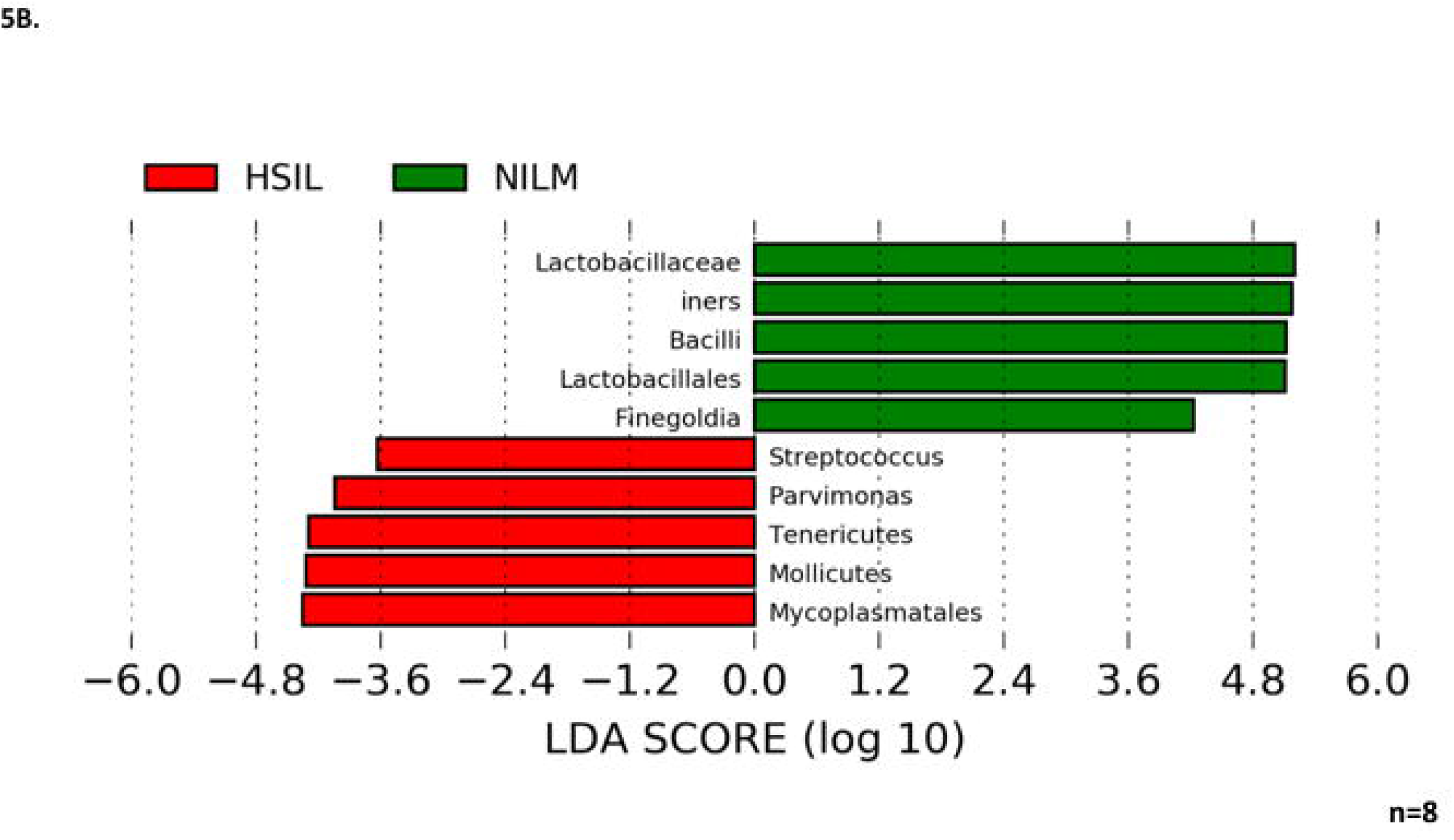
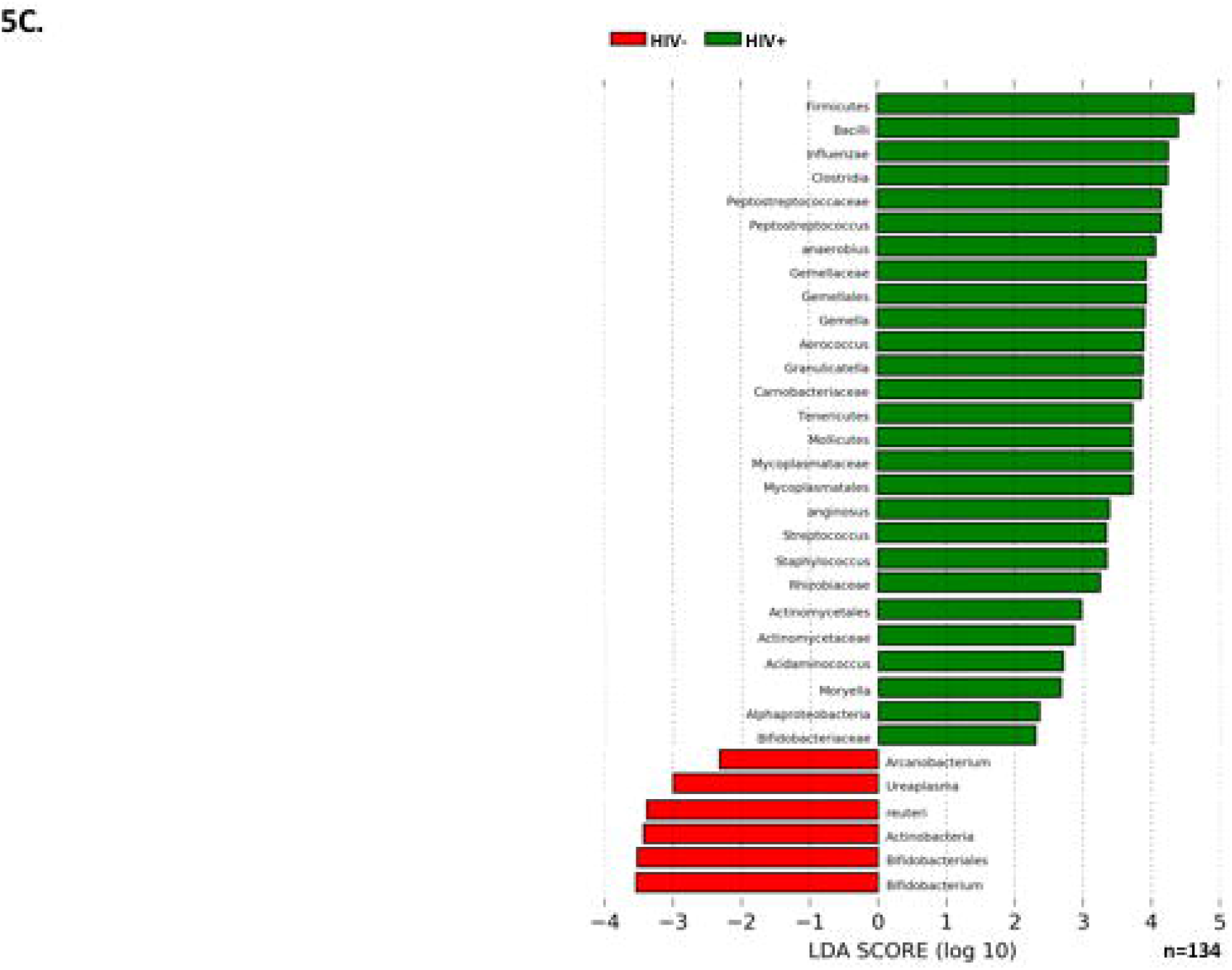
LEfSe LDA scores: Microbes associated with cervical cytology status and/or HIV status are displayed. **A**. Taxonomies differentiating bacterial microbiota in normal vs. HSIL cervices. **B**. Taxonomies differentiating bacterial microbiota in normal vs. HSIL cervices in age-matched HIV+ patients. **C**. Taxonomies differentiating bacterial microbiota in HIV− vs. HIV+ cervices.

## Discussion

It is well established that certain members of the cervicovaginal microbiome protect against infection and pathogenesis. The primary defense mechanisms of the cervicovaginal mucosa are antimicrobial peptides, a pH of less than 4.5, and a microbiome dominated by Lactobacilli. An imbalance in these defenses can result in physiochemical changes which produce alterations of the vaginal mucosa and cervical epithelium [28]. In particular, an abundance of *Lactobacillus crispatus* shows an inverse relationship with detectable or symptomatic HIV, HPV, or herpesvirus infection [29]. This suggests that other cervicovaginal microbes may be important in preventing or enhancing the acquisition and pathogenesis of such infections. Microbes which associate with enhanced pathogenesis have largely gone unidentified or unstudied, especially in the most at-risk population: HIV positive women in Sub-Saharan Africa.

In this study, HIV was shown to have a significant effect on the cervical microbiome, increasing bacterial richness and decreasing beta diversity. These results are similar to what has been reported on the cervicovaginal microbiome, and suggest changes in the cervical epithelium microenvironment brought on by HIV exert some selective pressure on cervical bacteria communities. Mycoplasma was significantly more abundant in HIV positive patients, and was found to be one of the main categories of bacteria which differentiate the cervical microbiota of HIV positive and HIV negative individuals. Interestingly, bacteria of the order Bacilli, of which Lactobacillus is a member, were strongly associated with HIV positive patients. The absence of Bacilli reads classified as Lactobacillus among the significant factors of HIV positive cervical microbiota suggests this may be due to a shift from protective to non-protective Bacilli in HIV+ individuals. When the cohort was analyzed without taking HIV status into account, cervical cytology did not appear to have a statistically significant association with differences in the cervical microbiome. However, when HIV was controlled for by separating analysis into groups of HIV positive or HIV negative only, differences in cervical bacterial communities which varied on the basis cervical lesion status began to reach statistical significance. This suggests developing pre-cancerous cervical lesions is associated with certain microbiota. Among these microbiota, Mycoplasmatales stood out as the most significant differentiator between the cervical microbiota of a cervix with pre-cancerous lesions from one without. Bacteria belonging to the family Mycoplasmatales also showed the clearest linear increase in abundance with development of more severe lesions in both HIV positive and HIV negative populations. The most common Mycoplasmatales to infect the urogenital tract of women is *Mycoplasma genitalium. M. genitalium* is a non-commensal bacteria commonly associated with other STIs, especially HIV [30–32]. The temporal relationship between HIV and *M. genitalium* infection is unclear. It is not well understood if HIV promotes mycoplasma initial infection, or persistence of what may have been a transient infection in an HIV-negative individual. One study found that HIV-positive women cleared *M. genitalium* infections more slowly in comparison with HIV− negative women, and the infection recurred in 39% patients after clearance [33]. The role of *M. genitalium* infection in influencing initial infection of HIV also remains unclear, however a strong association between severity of *M. genitalium* infection and HIV shedding from the cervix has been shown [34]. What is clear is that *M. genitalium* infects the epithelia, disrupting tight junctions, and inducing a chronic inflammatory response. The potential for *M. genitalium* to influence replication of HIV suggests that host innate responses to *M. genitalium* infection may influence pathogenesis of other sexually transmitted infections. Induction of HPV in this way is particularly interesting based on the association between Mycoplasma and cervical lesions. Infection with *M. genitalium* increases the rate of infection with an HPV genotype associated with a high-risk of developing cervical cancer [35]. Recent work has shown that mycoplasma also increases the risk of development of cervical lesions, supporting the association we report in this study [35]. Mycoplasma can establish persistent, intracellular infections in epithelia cells, which may lead to bacterial vaginosis and/or cervicitis. *M. genitalium* has been established as an independent, causal microbe responsible for cervicitis [36]. This suggests that Mycoplasma may act as both an intracellular and extracellular stressor, particularly if coinfection with HPV has taken place. This interaction would most likely involve inflammatory cytokines induced by Mycoplasma infection. Further study is needed to determine if the inflammatory cytokines induced by Mycoplasma infection include those which are associated with pre-cancerous cervical lesions.

Mycoplasma is a low-abundance microbe which has been shown to cause cervicitis. However, the lack of significant associations in previous metagenomic studies, is largely due to a lack of optimization of statistical analyses for the presence of low abundance microbes. In our study, Mycoplasma was a prominent result, likely due to the large HIV positive proportion of the cohort, wherein immunosuppression allowed higher abundance of the bacteria to accumulate. Statistical analysis of just the HIV negative portion of the cohort did not identify Mycoplasmatales as a significant factor. However, a linear increase in the abundance of Mycoplasmatales from NILM to HSIL seen in both HIV positive and negative groups.

Analysis of HIV negative, NILM samples showed an abundance of anaerobic bacteria. This may be a result of the sampling method used, rather than a representation of the average HIV negative NILM cervical microbiome in Tanzania. Because samples were only obtained from women attending the women’s clinic, it is possible that some of the women had slightly unusual microbiota due to bacterial infections, but were NILM by the pap smear. If this were the case, it would explain the high contribution of anaerobic and/or pathogenic bacteria in NILM women attending the clinic. Another possibility is that cervicovaginal infection with anaeroabic bacterial in the general population is higher than previoiusly thought.

In this study, we took great effort to control for variation in the cervical microbiome so as to reduce confounding effects that might obsure the bacterial communities which were associated with HPV pathogenesis. The HIV positive population is of particular interest, since they appear to show enhanced cervical microbiota associated with HPV pathegenesis. In future studies, recruiting a cohort of all HIV positive women with and without cervical lesions would be desirable in order to better characterize HIV-associated microbiota which promote HPV infection and progression to cervical cancer. Currently, no study has been conducted with such a focused and controlled group. It is clear that variables such as diet, genetic background, antibiotics or ART, can dramatically effect the microbiota, and thus should be carefuly controlled at the point of recruitment to the study.

Longitudinal studies of the cervical microbiome are needed to understand how microbe populations change over time, particularly in individuals with HSIL. Long-term longitudinal studies will be able to determine if changes in the cervical microbiota preempt and predict the development of lesions, or if the shift in microbiota happens after lesions have developed. Because progression of HPV infection to cervical cancer is a process that takes decades, and in many individuals never reaches cancer at all, such a study would need to be large. Studies of the cervical microbiome can be further improved using metagenomic sequencing, rather than 16s or other targeted sequencing techniques which lack depth. 16s amplification ignores microbes which lack a gene to match the primers, for example; viruses, archaea, and eukaryotes are not accounted for. Because only a portion of one gene is being sequenced, the microbes present may only be estimated at the genus level or worse. Since the majority of medium or large scale cervicovaginal microbiome studies have used this method, the role of non-bacterial components of cervicovaginal microbiome in HPV infection and disease has not been characterized.

As the world’s HIV positive population grows, cervical cancer is expected to become an even more significant problem, despite increasing coverage of anti-retroviral treatment (ART). Compared to the risk reduction after ART seen in other AIDS-defining cancers like Kaposi’s sarcoma and non-Hodgkin’s lymphoma, the risk of cervical cancer is not significantly affected, and recurrence rates remain high with or without treatment [37–40]. Studies in this area suggests that progression of HPV infection depends on immunological status of the host such that ART is only able to indirectly affect HPV pathogenesis, potentially through an effect on circulatory CD4+ cell count. Identifying which aspects of the local and systemic effects of HIV infection contribute to progression from latent HPV infection to cervical cancer is crucial to understanding and predicting HPV pathogenesis. Current knowledge suggests effects on the cervical immune microenvironment may be key in this process. Understanding microbes which influence this environment will help identify cervical microbiota associated with low and high-grade cervical lesions. This may allow certain cervical microbiota to be used as diagnostic markers for those at high risk of developing cervical cancer, and for the development of preventative probiotic or antibiotic treatments which could control the cervical microbiome by promoting bacterial colonization with a microbiota associated with healthy cervical cytology. Our studies have identified a unique microbiota associated with HSIL. Data derived from of our precise sampling of cervical lesions leads us to propose that Mycoplasma contributes to a cervical microbiome status which promotes HPV-related cervical lesions. These results suggest a greater influence of the bacterial microbiota on the outcome of HPV infection than previously thought.

## Supporting Information

**Figure S1**: Bacterial 16s deep sequencing data was analyzed with rarefaction curves generated from the OTU data. These rarefactions were then compared with HIV status and cervical cytology. **S1A**. The red square represents HIV negative: “0,” and the black square represents HIV positive: “1.” The line indicated as “NA” is the unadjusted control. **S1B**. The red square represents HSIL, and the blue square represents LSIL. The green square represents NILM. The line indicated as “NA” is the unadjusted control.

**Figure S2**: The bacterial community composition differences were analyzed in reference to cervical cytology and HIV status using db-RDA with the unweighted UniFrac distance matrix. **S2A** Db-RDA analysis of the bacterial community as a function of HIV status. **S2B**. Db-RDA analysis of the bacterial community as a function of HIV status. **S2C**. Db-RDA analysis of the bacterial community as a function of cytology (NILM vs, LSIL or HSIL) at p=0.849. **S2D**. Db-RDA analysis of the bacterial community as a function of cytology (NILM vs, LSIL or HSIL) at p=0.05.

## Acknowledgements

We thank the Angeletti and Wood laboratory members and members of the NCV for critical discussions of this work. We thank members of the ORCI and the Tanzanian Ministry of Health, who facilitated these studies. We thank Danielle Shea for logistical and technical support related to this project.

